# Oxytocinergic modulation of threat-specific amygdala sensitization in humans is critically mediated by serotonergic mechanisms

**DOI:** 10.1101/2020.05.20.105684

**Authors:** Congcong Liu, Chunmei Lan, Keshuang Li, Feng Zhou, Shuxia Yao, Lei Xu, Ning Yang, Xinqi Zhou, Jiaxin Yang, Xue Yong, Yina Ma, Dirk Scheele, Keith M. Kendrick, Benjamin Becker

## Abstract

**Background:** Overarching conceptualizations propose that the complex social-emotional effects of oxytocin (OXT) in humans are partly mediated by interactions with other neurotransmitter systems. Recent animal models suggest that the anxiolytic effects of OXT are critically mediated by the serotonin (5-HT) system, yet direct evidence in humans is lacking.

**Methods:** To determine the role of 5-HT in OXT-induced attenuation of amygdala threat reactivity and sensitization/ desensitization, we conducted a parallel-group randomized placebo-controlled double-blind experiment during which n = 121 healthy subjects underwent a transient decrease in 5-HT signaling via acute tryptophan depletion (ATD, TRYP^-^) or the corresponding placebo-control protocols before the administration of intranasal OXT or placebo intranasal spray, respectively. Mean and repetition-dependent changes in threat-specific amygdala reactivity towards threatening stimuli (angry faces) as assessed by fMRI served as the primary outcome.

**Results:** No treatment main or interaction effects on amygdala threat reactivity were observed, yet OXT switched bilateral amygdala threat sensitization to desensitization and this effect was significantly attenuated during decreased central 5-HT signaling via pretreatment with TRYP^-^.

**Conclusions:** The present findings provide the first evidence for a role of OXT in threat-specific amygdala desensitization in humans and suggest that these effects are critically mediated by the 5-HT system. OXT may have a therapeutic potential to facilitate amygdala desensitization and adjunct up-regulation of 5-HT neurotransmission may facilitate OXT’s anxiolytic potential.

The trial was preregistered on clinicaltrials.gov (https://clinicaltrials.gov/ct2/show/NCT03426176, ID NCT03426176)

## Introduction

The hypothalamic peptide oxytocin (OXT) regulates a broad range of peripheral and central functions (1). Across species, OXT plays an important role in complex social behavior and basal emotion processes, particularly salience and threat processing (2). Overarching conceptualizations of the role of OXT in human social-emotional behavior have proposed that the complex behavioral effects of OXT are partly mediated by interactions with other neurotransmitter systems (3). Such interactions have been evidenced by initial animal models demonstrating that OXT’s effects in the domains of pair bonding are partly mediated by dopamine (4) whereas social reward and anxiolytic effects involve interactions with the serotonin (5-HT) system (5–7).

Accumulating evidence further suggests that the anxiolytic properties of OXT are (partly) mediated by the 5-HT system. Direct evidence for a role of 5-HT in OXT’s anxiolytic effects has been demonstrated in a seminal rodent model combining genetic editing with OXT infusion (7). This study demonstrated that OXT receptors are expressed in one third of the 5-HT releasing neurons in the raphe nucleus, which represents the principal source of central 5-HT as well as afferent serotonergic projections to the amygdala (8). More recently, initial studies combined intranasal or intracerebroventricular OXT administration with concomitant molecular imaging of 5-HT neurotransmission in non-human primates and humans and reported OXT-induced modulations of serotonergic signaling in regions strongly engaged in salience and threat processing, particularly the amygdala and insula, with further analyses suggesting a central role of the amygdala in the oxytocinergic regulation of 5-HT release (6,9).

To directly examine whether the anxiolytic effects of OXT on threat-related amygdala reactivity in humans are mediated by the 5-HT system, we conducted a parallel-group randomized placebo-controlled double-blind fMRI experiment during which N=121 healthy male participants who either underwent transient decreases in 5-HT signaling (via acute tryptophan depletion, ATD; TRYP^-^) or a matched control protocol (TRYP^+^) before the administration of intranasal OXT or placebo (PLC). Based on previous animal models we specifically hypothesized that (1) OXT would dampen amygdala threat reactivity to angry faces relative to placebo, and that (2) pretreatment with 5-HT depletion would attenuate OXT-induced dampening of amygdala responses relative to OXT treatment alone.

Amygdala responses show rapid adaptations with repeated stimulus presentations with both reduced (desensitization) (10–12) and increased reactivity (sensitization) (13–15) being reported dependent upon the respective amygdala subregions involved and emotional content. Furthermore, a higher retest reliability of repetition-dependent amygdala signal changes was found when compared to mean amplitude measurements, suggesting a particular stable marker for pharmacological imaging (e.g. Gee et al., 2015). Although a number of previous studies suggests that amygdala sensitization/desensitization is modulated by serotonergic signaling (17) and that OXT may modulate arousal and amygdala habituation in trust and cooperation contexts (18,19), the role of OXT for threat-related habitation is still unclear.

Therefore, we additionally explored the interactive effects of OXT and 5-HT on sensitization/desensitization employing a comparable analysis strategy as for amygdala mean amplitudes. Given that previous rodent models demonstrated that OXT regulates amygdala threat-responses via direct hypothalamic-amygdala neuronal projections (20) as well as indirect pathways via OXT receptors expressed on serotonergic raphe neurons (7), we hypothesized that downregulation of serotonergic signaling would decrease but not fully abolish the effects of intranasal OXT on amygdala threat reactivity. Finally, although amygdala desensitization as assessed by the mean of a block difference approach represents a robust and comparably reliable within-subject fMRI index (21), we additionally included an independent dataset to determine the robustness of threat-specific amygdala sensitization/desensitization across samples.

## Methods and materials

### Participants

The primary aim of the study was to determine interactions between the OXT and 5-HT systems on threat-related amygdala reactivity and amygdala sensitization/desensitization in humans. A total of *N*=121 right-handed healthy male participants were enrolled in the present randomized placebo-controlled double-blind between-subject pharmacological fMRI study. To reduce variance related to sex differences in threat-related amygdala responses and the effects of oxytocin on amygdala reactivity (e.g. Gao et al., 2016) only male participants were included. Given the complexity of the design, a pragmatic approach for sample size determination was employed based on a recent fMRI study (23) comparing effects of different OXT dosages on threat-related amygdala activity (for a similar approach see recent study comparing OXT with another anxiolytic agent) (24). Exclusion criteria included: (1) current/history of physical or psychiatric disorders, (2) current/regular use of licit or illicit psychotropic substances, (3) weight >85 kilograms, (4) MRI contraindications, (5) cardiovascular disorders including high blood pressure, (6) contraindications for either OXT or acute tryptophan depletion (ATD) protocol. Based on initial quality assessments data from n=9 subjects were excluded from the analysis (n=5, technical problems during data acquisition; n=2, performance >3 SDs from mean accuracy suggesting a lack of attention or adherence to the experimental protocols; n=2, history of mania or depression details see CONSORT flowchart, **Figure S1**).

Study protocols were approved by the ethics committee (University of Electronic Science and Technology of China) and adhered to the latest revision of the Declaration of Helsinki. Written informed consent was obtained and the study was pre-registered on clinicaltrials.gov (https://clinicaltrials.gov/ct2/show/NCT03426176, ID NCT03426176).

### Procedure

The present study employed a between-subject randomized double-blind pharmacological fMRI design incorporating four treatment groups which received combinations of ATD (TRYP^-^ versus TRYP^+^ drink) and intranasal oxytocin (oxytocin versus placebo nasal spray). Participants were instructed to abstain from alcohol and caffeine for 24h and from food and drinks (except water) for 12h prior to the experiment. To adhere to the pharmacodynamic profile of the treatments participants arrived between 7:30 to 10:00AM and underwent fMRI acquisition between 13:30 to 16:00PM. Upon arrival, participants received a standardized protein-poor food for breakfast. Following the assessment of pre-treatment control variables participants were administered either a tryptophan-depleted amino acid (TRYP^-^) or a control drink (TRYP^+^) followed by a resting period of 5h to achieve a robust reduction in tryptophan levels. Subsequently, control variables were assessed and 5h after the amino acid drink participants administered either OXT (24IU) or placebo (PLC) nasal spray (standardized according to previous oxytocin administration studies) (25). In line with the pharmacokinetic profile of intranasal OXT(23), the fMRI paradigm was scheduled 50min after OXT administration. Control variables were assessed before and after fMRI acquisition (schematic outline of the experimental protocols see **Figure 1**)

**Figure 1.**
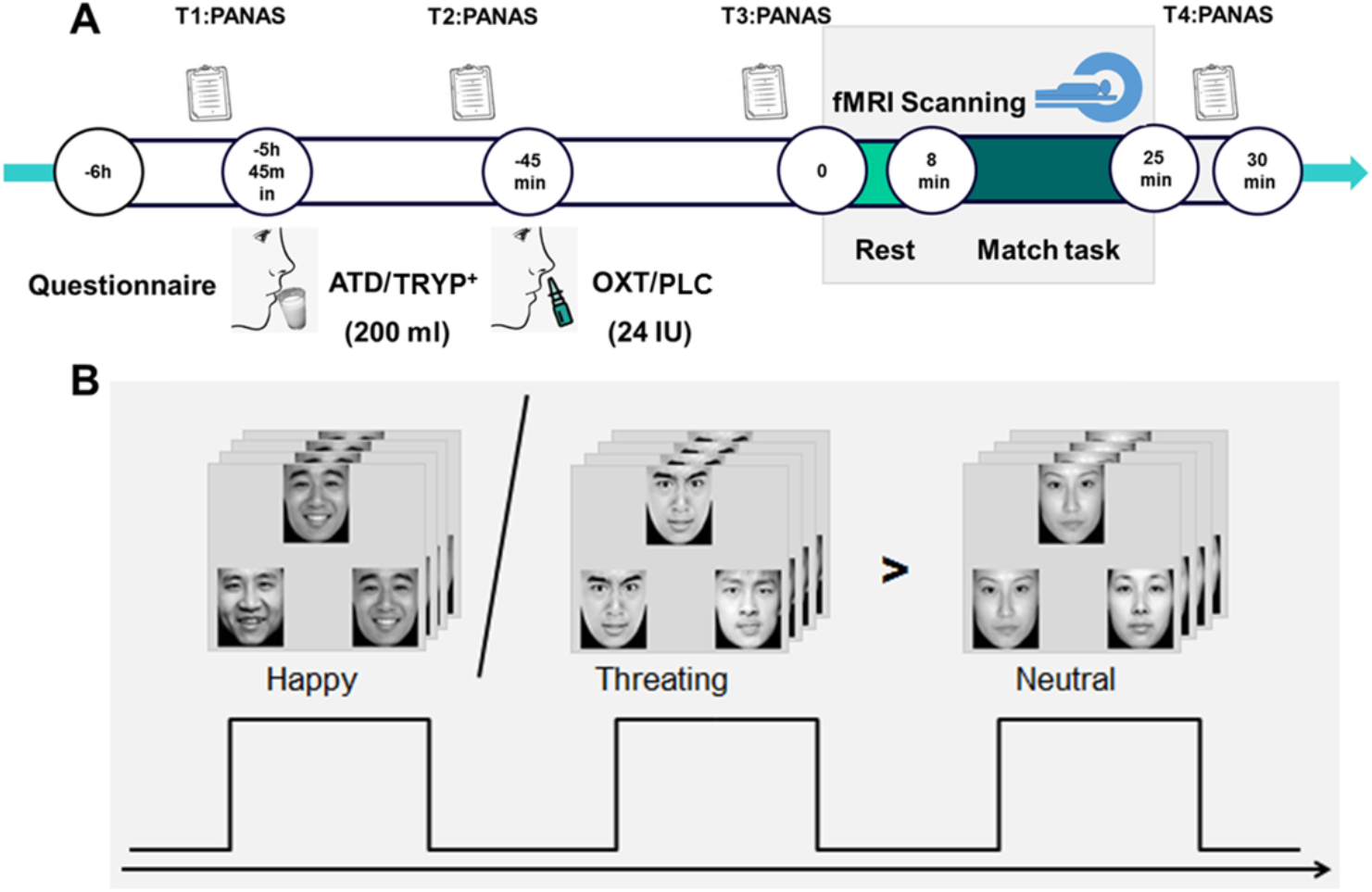
Experimental design and treatment protocols

### Control variables

To control for between-group differences in depressive symptom load, anxiety and current stress, the Beck Depression Inventory (BDI-II) (26), State-Trait Anxiety Inventory (STAI) (27) and Perceived Stress Scale (PSS) (28) were administered before treatment administration. To assess effects of treatment on mood during the entire experimental procedure, the Positive and Negative Affect Schedule (PANAS) (29) was repeatedly administered before administration of the amino acid drink (T1) and the nasal spray (T2) as well as immediately before MRI acquisition (T3) and at the end of the experiment (T4).

### Serotonin Dietary Manipulation (oral administration)

A previously validated dietary drink (TRYP^-^) inducing ATD and a control protocol (TRYP^+^) (30,31) was used to temporarily lower central 5-HT level (TRYP^-^ group) in a randomized, double-blind, placebo-controlled, between-subjects design (Supplementary Material).

### Oxytocin (intranasal administration)

Oxytocin (OXT) nasal spray comprised oxytocin, glycerine, sodium chloride and purified water, the placebo nasal spray (PLC) included identical ingredients except for oxytocin (both provided in identical spray bottles by Sichuan Meike Pharmaceutical Co. Ltd, Sichuan, China). In line with previous intranasal OXT administration studies (32) a single dose of 24 international units (IU) was administered with 3 puffs per nostril.

### Experimental paradigm

The blocked-design fMRI paradigm has been previously validated and demonstrated to produce robust amygdala responses in response to threatening (angry) faces (21,33). The paradigm consisted of 3 runs and every run comprised 6 blocks of facial stimuli as well as 2 blocks of non-facial stimuli serving as non-social control stimuli. During the face-processing blocks, a trio of condition-specific (neutral, angry or happy expressions) facial stimuli was presented and subjects required to select one of the two faces (bottom) that was identical to a target face (top). Each block comprised four condition-specific trials, balanced for gender.

Asian facial stimuli were selected from a standardized Asian facial expression database (34). During the non-social control blocks a trio of simple geometric shapes (circles and ellipses) was presented and subjects were required to select one of two shapes (bottom) that matched the target shape presented on the top (**Figure 1B)**. Each control block comprised four different shape trios. All blocks were preceded by a brief instruction (‘Face match’ or ‘Shapes match’) that lasted 2s. Within each block, each trial was presented for 4s with a variable interstimulus interval (ISI) of 1-3 s (mean, 2s). The total block length was 24s and the total paradigm lasted 16min 48s.

### MRI data acquisition and processing

MRI data was acquired on a 3 Tesla MRI system and preprocessed using routines in SPM 12 (see **Supplemental Material**). On the first level a general linear model (GLM) was employed and included condition-specific regressors modelling the experimental conditions, the cue-phase and the six head motion parameters. To examine (de-)sensitization effects, a separate first level model was designed which additionally modeled the blocks separately. The corresponding design matrices were convolved with the default SPM hemodynamic response function (HRF). The design matrices additionally included a high pass filter to control for low frequency components and a first-order autoregressive model (AR[1]) to account for autocorrelation in the time-series. To evaluate our a-priori hypotheses analyses focused on threat-specific brain activity using [angry > neutral faces] as the primary contrast of interest.

### Statistical Analysis

#### Effects on mean amygdala threat reactivity

Effects on threat-related amygdala reactivity were examined using a standard GLM approach employing the mean contrast of all angry facial expression blocks relative to neutral faces [angry_all-blocks_ > neutral_all-blocks_]. On the second level, effects of treatment were examined by means of mixed ANOVA models including treatments (amino acid mixture, TRYP^-^/TRYP^+^ and intranasal spray OXT/PLC) as between-subject factors.

#### Effects on amygdala threat sensitization/desensitization

Effects on amygdala threat sensitization/desensitization were analyzed using the mean of a block difference model including the first and last block (first block minus last block, FmL) which is more sensitive compared to the means of the regression approach with respect to complex non-linear dependencies during habituation (35). To this end, amplitude differences between the first block in the first run and the corresponding last block in the last run were calculated. To separate threat-specific amygdala habituation from unspecific habituation to facial stimuli (e.g. Fischer et al., 2003), the primary outcome employed a subtraction of the neutral facial stimuli [(angry_first-block_>neutral_first-block_) > (angry_last-block_>neutral_last-block_)]. These contrast images were subjected to second level mixed ANOVA models including amino acid mixture (TRYP^-^ vs. TRYP^+^) and intranasal spray (OXT vs. PLC) as between-subject factors.

#### A-priori region of interest and statistical thresholding

In line with our regional a-priori hypotheses, previous evidence that the anxiolytic effects of OXT in rodents and humans are mediated by the amygdala and a central role of the amygdala in OXT-induced 5-HT release in humans (9), analyses focused on the amygdala as a-priori region of interest. To this end, a bilateral mask for the entire left and right amygdalae were defined using the Automatic Anatomical Labelling (AAL) template (37) and employed for Family-Wise Error (FWE) correction using a small-volume correction (*p*_FWE_ < 0.05, svc). An additional exploratory whole-brain analysis was computed to explore treatment interaction effects in regions outside of the a-priori defined region of interest using a whole-brain threshold of *p*_FWE_ < 0.05. For post-hoc comparisons individual parameter estimates were extracted from the amygdala mask. To evaluate our hypotheses post-hoc comparisons focused on comparing the treatment groups with the respective PLC-treated reference group.

#### Robustness of threat-specific amygdala sensitization / desensitization

To further determine the robustness of threat-specific amygdala sensitization / desensitization we included data from an independent sample of n = 25 healthy males who underwent a control protocol (TRYP+) administration before fMRI acquisition with a similar block design paradigm (details see **Supplemental Material**). In line with the analysis in the original sample the second level analysis focused on the sensitization / desensitization contrast [(angry_first-block_>neutral_first-block_) > (angry_last-block_>neutral_last-block_)].

## Results

### Sample characteristics, confounders and mood

There were no pre-treatment group differences in age, depressive symptoms, anxiety, current stress levels and pre-treatment mood (all *p* > 0.16, details see **Table 1**). Examining effects of treatment on mood during the course of the experiment by means of mixed ANOVA models with amino acid mixture (TRYP^-^ vs. TRYP^+^) and intranasal spray (OXT vs. PLC) as between-subject factors and timepoint (T1-T4; pre-oral administration, pre-intranasal administration, pre-fMRI, post-fMRI, T1-T4) as within-subject factor revealed a significant main effect of time on both positive (*F*(3,306) = 20.03, *p* < 0.001, *η*^*2*^_*p*_ = 0.164) and negative affect (*F*(3,306) = 14.73, *p* < 0.001, *η*^*2*^_*p*_ = 0.126), suggesting a general decrease of mood over the experiment. Moreover, a significant interaction effect of ATD and OXT on negative affect (*F*(1,102) = 7.99, *p* = 0.006, *η*^*2*^_*p*_ = 0.073) was observed, with post-hoc analyses suggesting that when the participants received OXT treatment following TRYP^-^, they reported higher negative affect at T4 as compared to the PLC condition.

**Table 1.** Sample characteristics (n = 112)

|  | TRYP<sup>-</sup>-OXT<br>(n = 28) | TRYP<sup>-</sup>-PLC<br>(n = 26) | TRYP<sup>+</sup>-OXT<br>(n = 29) | TRYP<sup>+</sup>-PLC<br>(n = 29) | *p* Value |
| --- | --- | --- | --- | --- | --- |
| Age, Years | 21.89 ± 2.39 | 21.73 ± 2.66 | 22.24 ± 2.29 | 22.10 ± 2.13 | 0.86 |
| STAI-TAI | 41.79 ± 8.00 | 41.00 ± 5.85 | 38.72 ± 7.97 | 40.83 ± 7.00 | 0.43 |
| STAI-SAI | 38.77 ± 8.81 | 35.16 ± 6.36 | 34.71 ± 8.50 | 35.43 ± 6.64 | 0.21 |
| BDI | 6.07 ± 6.44 | 8.08 ± 6.05 | 4.90 ± 6.40 | 5.21 ± 4.97 | 0.21 |
| PSS | 14.50 ± 5.45 | 15.42 ± 5.25 | 13.59 ± 4.69 | 15.41 ± 4.58 | 0.46 |
| PANAS-P(T1) | 24.19 ± 7.17 | 24.08 ± 8.38 | 24.46 ± 6.95 | 26.04 ± 8.07 | 0.76 |
| PANAS-N(T1) | 15.88 ± 9.18 | 13.04 ± 6.09 | 11.75 ± 5.02 | 13.43 ± 5.95 | 0.16 |

Mean and standard deviations ( $\pm SD$ ) are displayed. Abbreviations: BDI, Beck's Depression Inventory; PANAS-N, Positive and Negative Affect Schedule-Negative affect; PANAS-P, Positive and Negative Affect Schedule-Positive affect; PSS, Perceived Stress Scale; STAI-TAI, State-Trait Anxiety Inventory-Trait Anxiety Inventory; T1, timepoint. T1, pre-treatment assessment.

### Behavioral results

Examining accuracy and response times by means of mixed ANOVAs with condition (angry face vs. happy face vs. neutral face vs. geometric shape) as within-subject factor and amino acid mixture (TRYP^-^ vs. TRYP^+^) and intranasal spray (OXT vs. PLC) as between-subject factors revealed no significant effects involving treatment on accuracy and reaction time (RT) arguing against potential confounding effects of treatment on basal attention and vigilance (details **Supplementary Material**).

### Effects on mean threat-related amygdala amplitude

Contrary to our hypothesis no significant main or interaction effects of ATD and OXT on amygdala threat-reactivity were observed (contrast [angry_all-block_>neutral_all-blocks_]).

### Effects on amygdala threat sensitization / desensitization

Examination of treatment effects on threat-specific amygdala sensitization / desensitization by means of voxel-wise ANOVA models including the between-subject factors treatment (amino acid mixture (TRYP^-^ vs. TRYP^+^) and intranasal spray (OXT vs. PLC) revealed a significant time (first block vs. last block) × treatment interaction effect in the bilateral amygdala (left (MNI [−21 −3 −15], *p*_FWE_ = 0.008, k = 20, *t* = 3.64; right, MNI [18 −3 −15], *p*_FWE_ = 0.019, k = 1, *t* = 3.25; small volume corrected for the bilateral amygdala mask) (**Figure 2**). Post-hoc comparisons on the extracted parameter estimates from amygdala mask (contrast of interest, angry_first-block_-neutral_first-block_ > angry_last-block_-neutral_last-block_) revealed that the placebo-treated reference group (TRYP^+^-PLC) demonstrated increased amygdala responses suggesting threat-specific sensitization rather than habituation of the amygdala. Compared to the reference group, OXT (TRYP^+^-OXT) switched amygdala sensitization to desensitization as reflected by significantly decreased threat-specific amygdala responses (*p*_FDR_ < 0.001; Cohen’s *d* = 0.99), while decreased serotonin signaling by TRYP^-^ pretreatment before OXT administration significantly attenuated this effect of OXT (TRYP^+^-OXT versus TRYP^-^ -OXT, *p*_FDR_ = 0.038; Cohen’s *d* = 0.52). An additional post hoc analysis on the condition-specific parameter estimates (neutral_first-block_ > neutral_last-block_ and angry_first-block_ > angry_last-block_, respectively) employing a two-way ANOVA with treatments as between subject factors aimed at further exploring whether the observed treatment effects were specifically driven by the angry face condition. A lack of significant effects on neutral faces in the context of a significant difference between the TRYP^+^-PLC group and TRYP^+^-OXT (*p*_FDR_ = 0.003; Cohen’s *d* = 0.83) and between the TRYP^+^-OXT and TRYP^-^-OXT (*p*_FDR_ = 0.037; Cohen’s *d* = 0.55) for the angry condition (angry_first-block_ > angry_last-block_) (**Figure 3**) further confirmed effects on threat-specific sensitization/desensitization.

**Figure 2.**
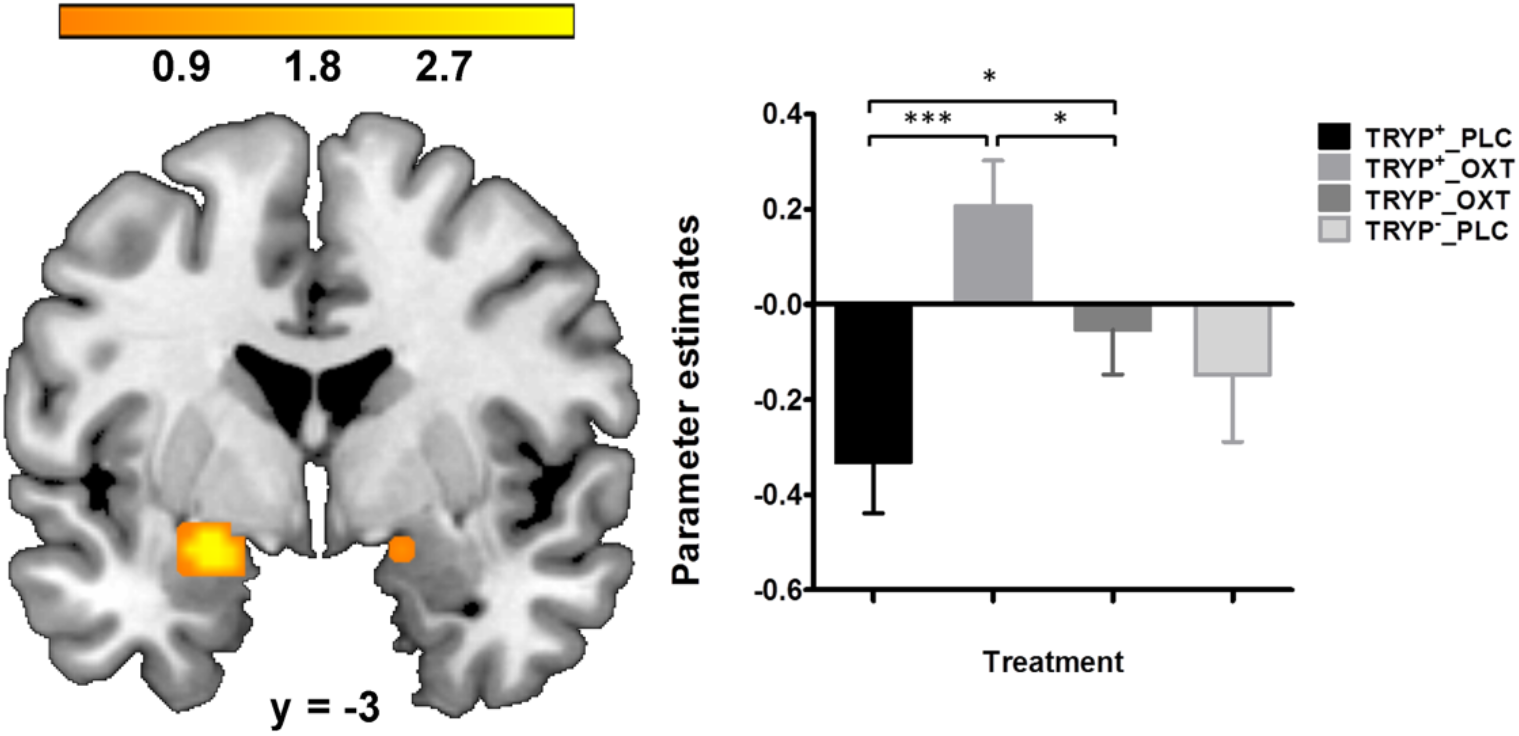
Effect of treatment on threat-specific amygdala sensitization/desensitization. The threat-specific effect in the bilateral amygdala is displayed at *p*_FWE-SVC_ < 0.05 thresholded for the entire bilateral amygdala. The color bar codes the t value. Bars on the right correspond to the extracted estimates for threat-specific sensitization/desensitization [(angry_first-block_>neutral_first-block_) > (angry_last-block_>neutral_last-block_)] for each treatment group. Results indicate that following placebo-treatment (TRYP^+^-PLC) the bilateral amygdala exhibited threat-sensitization, which was switched to desensitization following oxytocin (TRYP^+^-OXT) and that this effect of oxytocin was significantly attenuated, yet not fully abolished after pretreatment with acute tryptophan depletion (TRYP^-^-OXT). Abbreviations: TRYP^-^, acute tryptophan depletion; TRYP^+^, control treatment for acute tryptophan depletion; OXT, oxytocin nasal spray; PLC placebo for oxytocin nasal spray * and *** denote significant post-hoc differences at *p*_FDR_ < 0.05 and *p*_FDR_ < 0.001.

**Figure 3.**
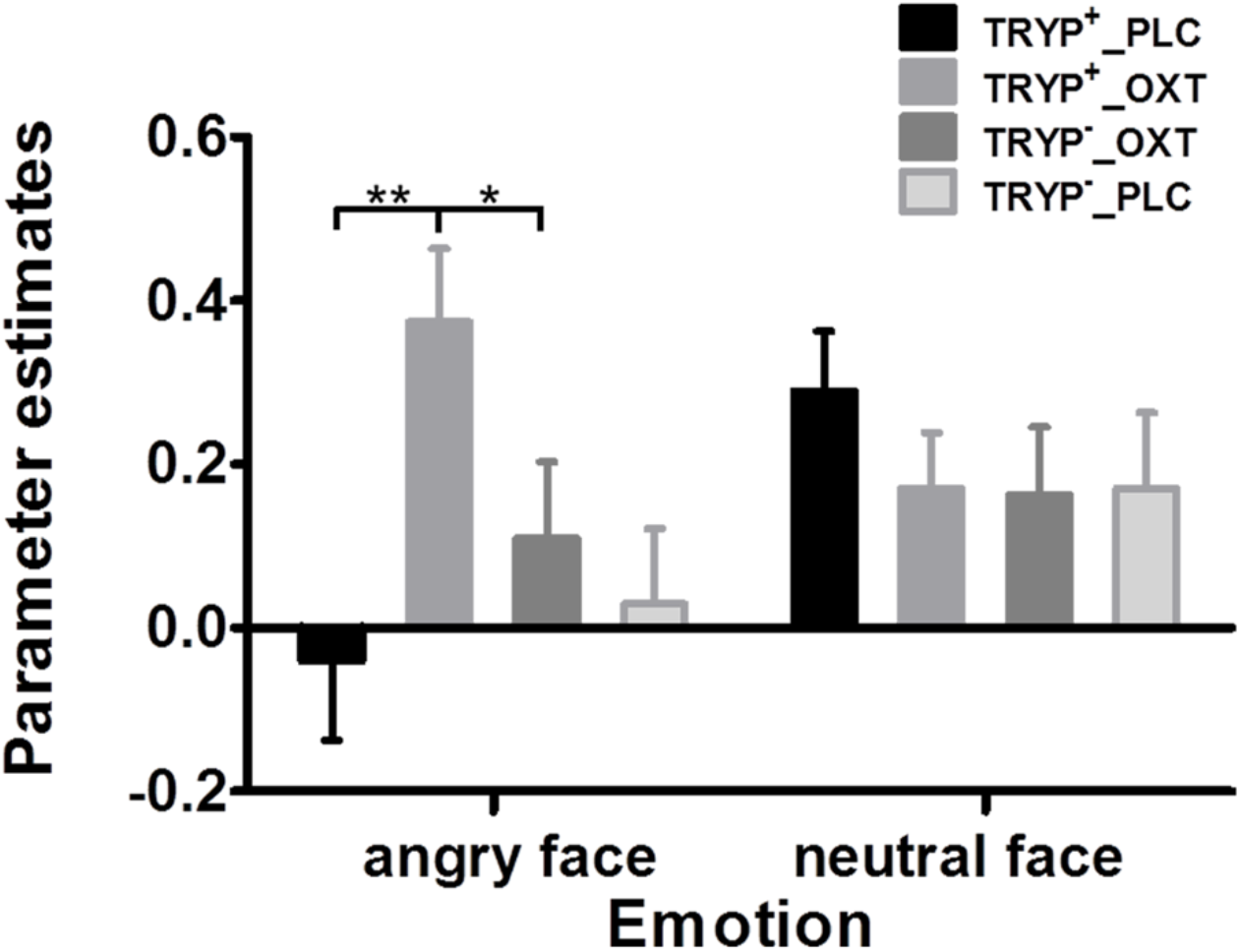
Condition-specific amygdala sensitization / desensitization Condition-specific parameter estimates for amygdala sensitization / desenitization for angry and neutral facees. Bars correspond to the extracted estimates for the identified amygdala region and suggest a threat-specific differences between the treatment groups. Abbreviations: TRYP^-^, acute tryptophan depletion; TRYP^+^, control treatment for acute tryptophan depletion; OXT, oxytocin nasal spray; PLC placebo for oxytocin intranasal spray * and ** denotes significant post-hoc differences at *p*_FDR_ < 0.01 and *p*_FDR_ < 0.05.

Additional control analysis examining effects of treatment on sensitization/ desensitization to positive stimuli (happy_first-block_-neutral_first-block_ > happy_last-block_-neutral_last-block_) and non-social stimuli (shapes_first-block_ > shapes_last-block_) did not reveal significant differences further suggesting threat-specific effects.

### Exploratory whole brain analysis

In line with our hypothesis the primary analysis focused on the amygdala. In addition, an unrestricted exploratory whole-brain analysis was conducted on the voxel-level. Results from this analysis revealed a significant time × treatment effect in cortical midline regions and the bilateral superior temporal gyrus. Post hoc analyses revealed a similar pattern as observed for the amygdala, specifically a threat-specific sensitization in the PLC group which was switched by OXT and attenuated following combined TRYP^-^ -OXT administration (details **Supplementary Material** and **Figure S2**).

### Robustness and replicability of threat-specific amygdala desensitization

Data from an independent validation sample of n = 25 healthy male subjects was employed to determine the robustness of the threat-specific amygdala sensitization observed in the control group (TRYP^+^-PLC). A voxel-wise second level analysis of the sensitization/desensitization contrast [(angry_first-block_>neutral_first-block_) > (angry_last-block_>neutral_last-block_)] confirmed threat-specific bilateral amygdala sensitization (*p*_FWE_ < 0.001) in the validation sample (**Supplementary Material** and **Figure S3**).

## Discussion

Overarching conceptualizations suggests a modulatory influence of OXT on 5-HT signaling and animal models demonstrated a functional relevance of this interaction for the anxiolytic potential of OXT. Building on these previous findings, the present pharmacological fMRI study employed an experimental protocol to reduce central 5-HT signaling before the administration of intranasal OXT to determine the role of 5-HT in mediating OXT-induced attenuation of amygdala threat reactivity. In contrast to our hypothesis no effects on the mean amplitude of amygdala threat reactivity were observed, however, further analyses on repetition-dependent threat-related amygdala reactivity revealed a sensitization of the bilateral amygdala with repeated presentation of threatening faces following control treatment (TRYP^+^-PLC), which was switched to desensitization following OXT (TRYP^+^-OXT) and that this effect of OXT was attenuated following decreased central serotonin signaling via pretreatment with TRYP^-^ (TRYP^-^-PLC). Together, these findings provide the first evidence that OXT facilitates amygdala threat desensitization and that this effect is (partly) mediated via a 5-HT dependent mechanism.

In contrast to our hypothesis, no effects of OXT on the mean amplitude of amygdala threat reactivity were observed which might be related to the specific threat stimuli chosen. We chose angry faces as direct threat stimuli and while previous studies demonstrated convergent evidence for a serotonergic modulation of amygdala reactivity to angry as well as fearful faces (38), the effects of intranasal OXT on amygdala threat-processing appear to depend strongly on the specific emotion of the faces displayed. Whereas previous intranasal OXT studies reported enhanced recognition of fearful faces and attenuated amygdala reactivity towards fearful faces, OXT did not consistently modulate recognition of or amygdala reactivity towards angry facial expressions (39–41), The differences may be explained in the different motivational tendencies inherent to the facial expressions, such that the dominant response to angry expressions is threat avoidance whereas the dominant response to fearful expressions is approach (42). Likewise, the observation that OXT reduces amygdala habituation to unreciprocated cooperation in men (19) indicates that the peptide’s effects on habituation may be domain-specific.

The amygdala exhibits rapid adaptations to repeated presentation of salient stimuli, including facial expressions (10) and these changes might be a more reliable marker of amygdala function as assessed by fMRI activation (16,21). Desensitization (habituation) of amygdala responses has been most consistently reported, but amygdala sensitization may also occur with repeated presentation of particularly salient or aversive stimuli (43). The currently prevailing dual-process framework proposes that the incremental (sensitization) and decremental (habituation) adaptations on the physiological and affective level are based on independent yet interacting processes (44). Sensitization has most consistently been observed in response to repeated presentations of reward- and threat-related stimuli (44) and sustained attention and less habituation to threat, including angry facial stimuli, has been demonstrated on the behavioral level (15,45). Consistently, increased neural responses with repeated presentation of angry emotional stimuli have been reported in limbic regions (15), including the amygdala (14). Partly resembling these previous observations, we found threat-specific amygdala sensitization in the control group (TRYP^+^-PLC) while OXT switched the direction of the repetition-dependent adaptation leading to a threat-specific habituation in this region.

The amygdala is particularly sensitive to social information (46) and exhibits widespread functional interactions with limbic and prefrontal systems (e.g. Bickart et al., 2014). Previous animal studies have pinpointed anxiolytic effects of OXT to the amygdala (20,48), and in humans intranasal OXT enhanced intrinsic communication between the amygdala and prefrontal regulatory regions (49). By modulating anger-related habituation of amygdala reactivity, OXT may facilitate rapid and flexible adaptation to social threat signals (3,49).

In line with a previous rodent model demonstrating that the anxiolytic effects of OXT are critically mediated by the 5-HT system (7), we found that TRYP^-^-induced reduction in serotonergic signaling attenuated, but did not fully abolished, the effects of OXT on amygdala threat reactivity. Animal models suggest that the anxiolytic action of OXT is mediated via hypothalamic-amygdala projection neurons (20) as well as OXT-sensitive receptors expressed on serotonergic raphe neurons (7). Whereas initial data suggests that TRYP^-^ may reduce peripheral OXT levels in the absence of behavioral effects (50), the combination of TRYP^-^ with placebo nasal spray did not produce effects on amygdala reactivity in the present study arguing against an TRYP^-^-induced unspecific decrease in OXT signaling. On the other hand, molecular imaging studies have demonstrated that intranasal OXT induces central serotonin release (6,9) and TRYP^-^ leads to stable and selective reductions in central 5-HT signaling (51), including attenuation of stimulated serotonin release (51,52) and availability of serotonin in presynaptic neurons (53). This suggests that pretreatment with TRYP^-^ diminished OXT-induced serotonin release via OXT-sensitive receptors on serotonergic raphe neurons in which in turn attenuated anxiolytic effects mediated by serotonergic raphe-amygdala pathways.

Given the increasing interest in the clinical application of OXT to attenuate anxiety and exaggerated amygdala responses (2,39), the present results have important clinical implications. First, deficient amygdala threat desensitization has been reported in several psychiatric disorders including anxiety disorders and autism and may represent a core pathological mechanism for the development and maintenance of exaggerated anxious arousal (17,54) with the present results indicating that OXT may facilitate amygdala threat habituation. Second, serotonin dysfunction is a core biomarker of anxiety and autism spectrum disorders (55,56). We found that the strength of the effect of OXT on amygdala threat desensitization was mediated by endogenous 5-HT levels, suggesting that individuals with low endogenous serotonergic levels may not fully capitalize on the anxiolytic effects of OXT and thus combined up-regulation of 5HT and oxytocin transmission may be needed for optimal facilitation of anxiolytic effects.

Findings of the present study need to be interpreted in the context of the following limitations. First, only male subjects were investigated due sex differences in 5-HT synthesis rate (57) and the effects of oxytocin on amygdala reactivity (19,22) and future studies need to determine whether the observed effects generalize to women. Second, tryptophan levels were not assessed in the present study, however, the study adhered to previously validated ATD protocols which have been shown to induce robust and selective decreases in 5-HT signaling (30,58). Third, OXT blood level were not assessed to validate the increase in (peripheral) OT levels, however, previous studies using identical intranasal administration protocols reported increased OXT blood levels (59,60). Finally, the present study did not include fearful facial expression stimuli. Although angry and fearful facial expressions are similar in valence and arousal (unpleasant and highly arousing), previous studies suggest different responses to angry and fearful facial expressions (42). Thus, these findings cannot be extrapolated to approach-related threat signals (e.g. fearful face).

## Acknowledgements

This work was supported by the National Key Research and Development Program of China (2018YFA0701400); National Natural Science Foundation of China (91632117, 31530032, 31700998); Open Research Fund of the State Key Laboratory of Cognitive Neuroscience and Learning, Beijing Normal University; Science, Innovation and Technology Department of the Sichuan Province (2018JY0001). The authors would like to thank all subjects who participate in this study.

