## supplements for "Oxytocinergic modulation of threat-specific amygdala sensitization in humans is critically mediated by serotonergic mechanisms"

**Authors:** Liu et al.

### Supplementary methods

#### Serotonin manipulation procedure

Previous studies have demonstrated that after administration of ATD tryptophan levels continuously decrease until it reaches a plateau after about 5 hours and that the robust decrease lasts around two hours (1-3). Given that tryptophan is the amino acid precursor of serotonin, the ATD procedure induces a transient selective reduction in central serotonergic neurotransmission (4). The contents of the amino-acid mixtures were based on previously validated proportions (3, 5, 6). The amino acid mixture (ATD, $TRYP^-$) consisted of 4.1g L-alanine, 3.7 g L-arginine, 2 g L-cystine, 2.4 g glycine, 2.4g L-histidine, 6g L-isoleucine, 10.1 g L-leucine, 6.7g L-lysine, 2.3 g L-methionine, 9.2g L-proline, 4.3 g L-phenylalanine, 5.2 g L-serine, 4.9 g L-threonine, 5.2 g L-tyrosine, 6.7 g L-valine, (total: 75.2g). The ATD control drink ($TRYP^+$) contained identical ingredients plus 3.0g of L-tryptophan (total: 78.2g). The drinks were prepared by stirring the mixture into 200-ml water and lemon-lime flavor was added to mask the taste of mixture.

#### MRI data acquisition and pre-processing

MRI data were acquired on a 3.0 Tesla GE MR750 system (General Electric Medical System, Milwaukee, WI, USA). T1-weighted high-resolution anatomical images were acquired with a spoiled gradient echo pulse sequence, repetition time (TR) = 5.9 ms, echo time (TE) = minimum, flip angle = 9°, field of view (FOV) = 256 × 256 mm, acquisition matrix = 256 × 256, thickness = 1 mm, number of slice = 156. For the functional MRI timeseries a total of 504 functional volumes were acquired using a T2*-weighted Echo Planar Imaging (EPI) sequence (TR = 2000 ms, TE = 30 ms, FOV = 220 × 220 mm, flip angle = 90°, image matrix = 64 × 64, thickness/gap = 3.2/0mm, 43 axial slices with an interleaved ascending order). Functional time-series were pre-processed using statistical parametric mapping (SPM12; Wellcome Department of Cognitive Neurology, Institute of Neurology, London, United Kingdom). For each subject and run the first seven volumes were discarded to allow for T1 equilibration. The remaining functional images were slice-time corrected and realigned to the first image to correct for head motion. The EPI images were next co-registered to the T1-weighted structural images and normalized to Montreal Neurological Institute (MNI) standard space using the segmentation parameters obtained from segmenting the structural images and interpolated at 3×3×3mm voxel size. Finally, the normalized images were spatially smoothed with an 8-mm full-width at half maximum (FWHM) Gaussian filter.

#### Robustness of threat-specific amygdala sensitization / desensitization in an independent sample

Using an identical fMRI paradigm a previous study demonstrated the robustness and within-subject replicability of amygdala desensitization as assessed by the mean of the block difference approach (7). However, the previous study employed a within-subject validation approach and examined amygdala habituation irrespective of emotional content of the faces whereas the present study examined threat-specific amygdala adaptations in a between-subject treatment design. To further test the replicability of the threat-specific amygdala sensitization / desensitization effects, we therefore employed a dataset from an independent study from our lab (details of this study see also pre-registration https://clinicaltrials.gov/ct2/show/NCT03549182, ID NCT03549182), henceforward referred to a 'validation study' in the context of the present manuscript). Briefly, in the validation study we employed the same paradigm, except that additionally a fearful face condition was included. Subjects were administered either an ATD or an ATD-placebo drink before fMRI acquisition, for the validation analysis the data from the n = 25 ATD-placebo treated healthy male subjects (mean age 21.44 ± 2.50) was included. The paradigm consisted of 6 runs and every run comprised 4 blocks of facial stimuli as well as 1 block of non-facial stimuli serving as non-social control stimuli. During the face-processing blocks, a trio of condition-specific (neutral, angry, fear or happy expressions) facial stimuli was presented and subjects required to select one of the two faces (bottom) that was identical to a target face (top). Each block comprised four condition-specific trials, balanced for gender. Asian facial stimuli were selected from a standardized Asian facial expression database (8). During the non-social control blocks a trio of simple geometric shapes (circles and ellipses) was presented and subjects required to select one of two shapes (bottom) that was identical to a target shape (top). Each control block comprised four different shape trios. All blocks were preceded by a brief instruction ('Face match' or 'Shape match') that lasted 2s. Within each block, each trial was presented for 4s with a variable interstimulus interval (ISI) of 1-3 s (mean, 2s). The total block length was 26s and the total paradigm lasted 21min 36s. Preprocessing and first-level modelling were conducted in SPM 12 and were identical to the procedures in the main study. In line with the original study, the main contrast of interest to determine the robustness of the amygdala sensitization / desensitization in the validation sample was $[(angry_{first\text{-}block} > neutral_{first\text{-}block}) > (angry_{last\text{-}block} > neutral_{last\text{-}block})]$.

### Supplementary results

#### Behavioral results

Significant main effects of condition were observed on both behavioral indices (accuracy: $F(3,324) = 5.69$, $p = 0.001$, $\eta^2_p = 0.051$; RT: $F(3,324) = 188.70$, $p < 0.001$, $\eta^2_p = 0.638$) with post-hoc analyses suggesting that accuracy for angry faces was significantly higher compared to all other conditions ($p$s < 0.001, all accuracies higher than 95%). Response times for geometric shapes (1045.34 ± 193.63) were faster compared to angry faces (1205.03 ± 230.79), angry faces compared happy faces (1279.80 ± 274.64) and slower response times for neutral faces (1351.16 ± 287.85) compared to all other conditions (all *p*s < 0.001).

**Exploratory whole-brain analysis**

The exploratory voxel-wise whole-brain analysis additionally revealed a significant sensitization ×treatment interaction effect in cortical midline regions (CMR), including right paracentral lobule, middle cingulate gyrus and precuneus (MNI [0 -12 51], $p_{FWE\text{-}cluster} = 0.014$, k = 185, t = 4.38), right superior temporal gyrus (STG, MNI [51 -18 6], $p_{FWE\text{-}cluster} = 0.002$, k = 283, t = 4.77), left superior temporal gyrus (MNI [-42 -24 3], $p_{FWE\text{-}cluster} = 0.006$, k = 229, t = 4.87) and right insula/superior temporal gyrus (MNI [39 -9 -9], $p_{FWE\text{-}cluster} = 0.032$, k = 147, t = 4.35, **Figure S2**). For visualization, the parameter estimates were extracted from a 6-mm sphere centered at the peak coordinates of the cluster (contrast of interest, $angry_{first\text{-}block}$-$neutral_{first\text{-}block}$ > $angry_{last\text{-}block}$-$neutral_{last\text{-}block}$) revealed that the reference group ($TRYP^+$-PLC) demonstrated a threat-specific sensitization effect in these regions, which was attenuated by oxytocin treatment ($TRYP^+$-PLC vs. $TRYP^+$-OXT, for bilateral STG, $p_{FDR} < 0.001$; for CMR, $p_{FDR} = 0.003$). However, following pre-treatment with ATD the effects of OXT were strongly attenuated as reflected by no significant differences between the $TRYP^-$-OXT group as compared to the $TRYP^+$-PLC group (for bilateral STG, $p = 0.158$; for CMR, $p = 0.266$), whereas there were significant differences between the $TRYP^-$-OXT and the $TRYP^+$-OXT groups (for bilateral STG, $p_{FDR} = 0.006$; for CMR, $p_{FDR} = 0.039$).

**Validation dataset – behavioral results**

Mixed ANOVAs with condition (angry face vs. happy face vs. neutral face vs. geometric shape) as within-subject factor and group ($TRYP^+$-PLC group from the current study vs. $TRYP^+$ group from validation study) as between-subject factor revealed no significant main effect of group and condition on accuracy and no significant main effect of group on reaction time (RT) except for a significant main effect of condition on RT (RT: $F(3,153) = 90.10$, $p < 0.001$, $\eta^2_p = 0.639$). Post-hoc analysis suggested that the response times for geometric shapes (1012.41 ± 199.75) were faster compared to angry faces (1164.12 ± 219.50), angry faces compared happy faces (1231.80 ± 276.55) and slower response times for neutral faces (1320.77 ± 290.00) compared to all other conditions (all *p*s < 0.001).

**Validation dataset – threat-specific amygdala sensitization / desensitization**

A voxel-wise one sample t-test on the validation dataset revealed a significant threat-specific sensitization effect in the right amygdala (MNI [27 3 -27], $p_{FWE} < 0.001$, k = 22, $t = 14.69$, small volume corrected for the entire bilateral amygdala) and left amygdala (MNI [-24 -9 -12], $p_{FWE} < 0.001$, k = 54, $t = 15.40$, small volume corrected for the entire bilateral amygdala) (**Figure S3**).

### Supplementary figures

**Figure S1** CONSORT flowchart

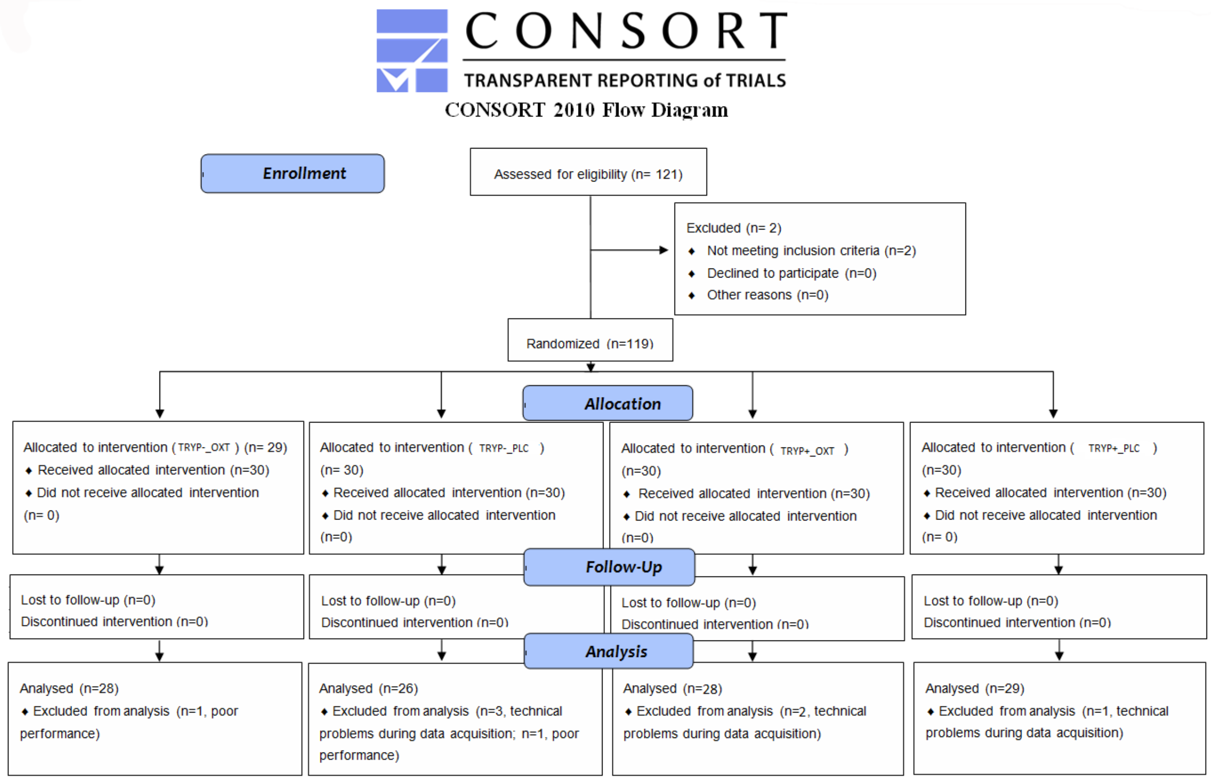

**Figure S2** Results from the exploratory whole-brain analysis of sensitization / desensitization differences between the treatment groups.

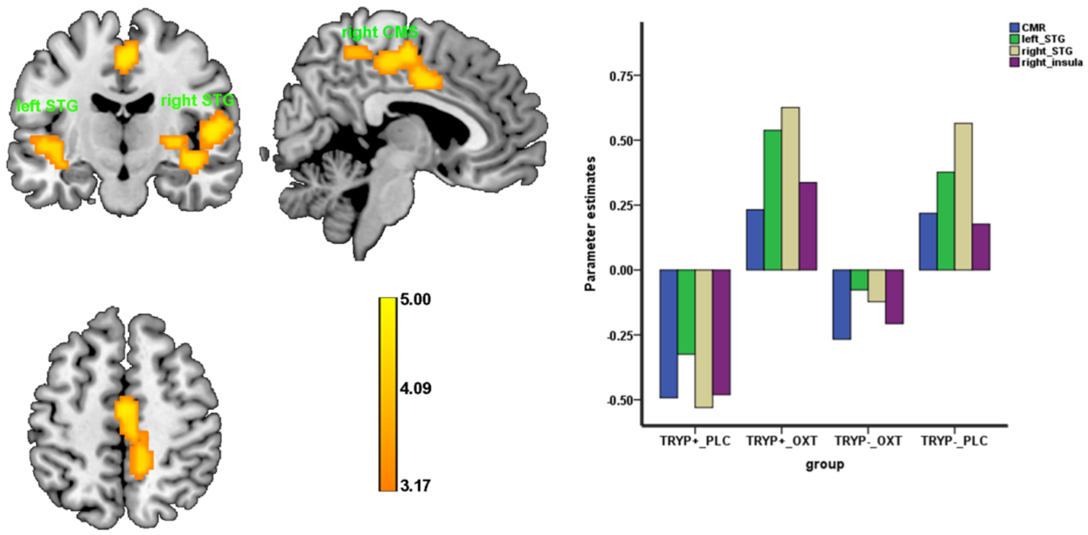


The exploratory whole brain analysis revealed significant time × treatment interaction effects on the whole brain level ($p_{\text{FWE - cluster}} < 0.05$).

Abbreviations: STG, superior temporal gyrus; CMR, cortical middle region, including right paracentral lobule/middle cingulum gyrus/precuneus.

**Figure S3** Results from the validation dataset

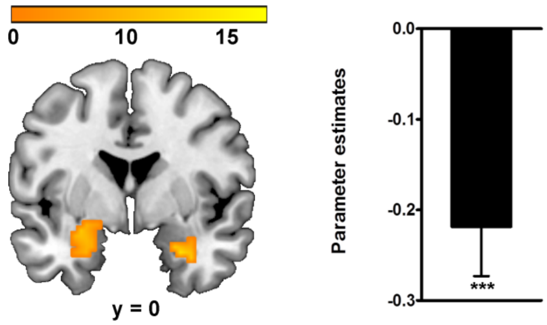


Examination of threat-specific amygdala sensitization / desensitization in the independent validation sample confirmed the robustness of increased amygdala reactivity over the course of the experiment (small volume corrceted for the entire amygdala, $p_{\mathrm{FWE}} < 0.001$). Extraction of parameter estimates further confirmed the threat-specific sensitization effects [(angry$_{\text{first-block}}$>neutral$_{\text{first-block}}$) > (angry$_{\text{last-block}}$>neutral$_{\text{last-block}}$). *** $p < 0.001$
